# A simplified analytical model of sound propagation in the human ear

**DOI:** 10.1101/571034

**Authors:** Wiktor L. Gambin

## Abstract

A simple mechanical model of sound propagation in the human ear from the entrance to the ear canal up to the round window membrane is outlined. The model shows the outer, middle and inner ear as two waveguides connected by a lever mechanism. The case when a sound wave from a sound source at a given frequency and intensity goes into the ear is considered. The sound as a plane elastic wave in the air of the ear canal is partially reflected from the eardrum and after relocation by a lever of the ossicles; it runs as a plane elastic wave in the cochlea fluid to be finally damped at the round window membrane. The basilar membrane excited by the running sound wave in the cochlea is taken as a chain of separate fibers. The power of the sound reaching the ear is compared with the power of the sound carriers in the ear. As a result, simple rules for the amplitude of the stapes footplate as well as for the amplitude and pressure of the forced acoustic wave in the cochlea are obtained. The formulas for the amplitudes of the membrane of the round window and the basilar membrane are also shown. The results of calculations based on these rules were compared with the measurements made on temporal bone specimens. The tests were done for the level of the sound source intensity of 90 dB and a set of frequencies from the range of 400-10,000 Hz. The amplitudes of the stapes-footplate and the round window membrane were measured. A mean difference, between the calculated and the measured values, for the stapes-footplate reached 31%, and for the round window membrane, it was 21%. A ratio of the basilar membrane velocity to the stapes footplate velocity as a function of the frequencies was shown. The obtained graph was close to that got by others as a result of the measurements.

## 1. Introduction

Over the past years, the problem of the perception of sound by the human ear enjoys great interest of researchers [1]. In order to properly describe the process of ***sound perception***, the sound wave pressure acting on the ear sound receptors must first be determined. To do it, one needs to have the right, but relatively simple, model of ***sound propagation*** in the human ear [2]. In this work is considered the case when a sound wave from a sound source at a given frequency and intensity goes into the ear. One can find the numerical models that study this problem in detail [3-5]. But our goal is to give some simple analytical rules for the amplitude of the sound pressure and amplitude of the sound wave running in the ear.

For this purpose the human ear is imagined as built of two waveguides connected by a mechanical set. The sound is transmitted by a plane elastic wave in air of the ear canal and then by the lever ossicles. After passing through the ear canal and reaching the tympanic membrane, there is a partial reflection of the sound wave. Part of the energy absorbed by the ossicles causes their vibrations, which in turn generate a sound wave propagating in the cochlea fluid. The sound wave flows around the basilar membrane, causing its waving, and finally is damped on the membrane of the round window.

To describe this process, the ***sound power transfer*** in the human ear is analyzed [6-7]. The sound power, reaching the ear in a unit of time is compared with the power of sound carriers in the ear. The air in the ear canal, the ossicles with the stapes footplate and the fluid in the cochlea are taken as the successive transporters of the sound energy. Some of the sound energy is absorbed by the basilar membrane. The last one is assumed to be a chain of independently vibrating fibers. The energy transfer ends at the round window membrane, which absorbs the energy of the running wave. One can determine parts of the sound energy transferred by each of these transporters. The sum of the determined parts of the sound energy must be equal to the input energy. In that way, the all parameters which describe the sound propagation in the ear can be shown as functions of the sound intensity level and the sound frequency that comes to the ear.

To validate the model, some of the got results were compared with the results of measurements. The values of the stapes footplate amplitudes, based on the received rule were compared with the measurements made on temporal bone specimens [8]. The tests were done for 90 dB and a set of fifteen frequencies from the range 400-10000 Hz. A mean difference, between the calculated and the measured values did not exceed 31%. The round window amplitudes were compared with the test results done for 90 dB and a set of four frequencies from the range 1000-8000 Hz [9]. A mean difference, between the calculated and the measured values did not exceed 18%. At the end, a ratio of the basilar membrane velocity to the stapes footplate velocity as a function of the frequencies was found. The obtained graph was close to the result of the measurements shown in [10] by Stenfelt et al.

## 2. Method

### Assumptions

To form a model of the outlined process, we introduce the following assumptions.

1. The outer ear is treated as a straight-line waveguide filled with air. The length of the waveguide is the length of the ear canal. The area of the waveguide cross-section change along its length. In the waveguide runs an ***elastic plane*** wave at the given level of sound intensity ***β*** and frequency ***f***_(***i***)_. The waveguide ends with an ***elastic*** tympanic membrane from which the sound wave is partially reflected.
2. The reduction of the sound intensity caused by the sound wave reflection is due to only the difference between the wave resistance of the air ***Z***_***0***_ in the ear canal and the wave resistance of the fluid ***Z***_***1***_ in the cochlea.
3. The middle ear is a lever system formed by a set of flexibly joined, stiff ossicles ended with a stapes footplate. The last one is suspended on a ***viscoelastic*** membrane of the annular ligament [11].
4. The inner ear is treated as a waveguide of changing cross-section filled with fluid, in which runs an ***elastic plane wave*** forced by vibrations of the stapes footplate. Its length **2*L***_***C***_ is a double distance from the oval window to the cochlea apex measured along the spiral axis of the cochlea. The axis of this waveguide is straight, but at the middle of the waveguide it changes direction to the opposite one. In this way, the waveguide is divided on two parallel connected parts, but which direct the sound wave in the opposite directions. The both parts of the waveguide are symmetric in relation to its center. There are no wave reflections on the transition between two parts of the waveguide [12].
5. Between two parts of the last waveguide lays a chain of ***uncoupled fibers*** of length ***L***_***BM***_ which replaces the basilar membrane. These ***viscoelastic*** fibers vibrate under the pressure of the sound wave running in the cochlea. The fibers have variable stiffness, high at the beginning of the chain and much smaller at its end. Because the stiffness and viscosity of the fibers are known, then one can find their resonant frequencies [13-14].
6. A waving of the chain of the fibers has no influence on the amplitude of the running sound wave. It will be shown that this effect really can be neglected. A justification for this assumption is given in the Appendix B. This fact is confirmed also by a numerical modelling of the process [12].
7. The other end of the above waveguide is closed by an ***elastic*** membrane of the round window, which absorbs all the energy of the running sound wave [15].

### Consequences

1. The damping of the sound wave amplitude in the cochlear fluid with respect to the amplitude of the wave in the ear canal is caused only by two factors. The first is the ***partial reflection*** of the wave from the tympanic membrane, and the second is the ***viscosity*** of the annular ligament. All other factors, such as wave reflections from the cochlea walls, basilar membrane waving, or flow through the basilar membrane opening are neglected.
2. The local increase or decrease of the sound wave amplitude in the cochlear fluid is caused only by the change in the cross-sectional area of the cochlea. As it travels, the wave grows up in amplitude, reaching a maximum at the cochlea apex, and then it goes down, when it comes on to the round window membrane.
3. A waving of the basilar membrane is caused by a difference in the sound wave pressures which take place in two parts of the waveguide separated by this membrane [12].

### Proceeding

A study of the sound propagation in the ***outer*** and ***middle ear*** gives the amplitude of the stapes footplate vibrations ***A***_***SF***_. At first, we calculate the power of the sound wave at the entrance to the ear canal, which allow us to find amplitude of the pressure acting on the tympanic membrane. An analysis of partial reflection of the sound wave and some its reinforcement due to the ossicular lever gives us the force ***N***_***SF***_ acting on the stapes footplate. A value of this force shows amplitude ***w***_***SF***_ of static displacements of the stapes footplate. Taking ***N***_***SF***_ as amplitude of the force which causes the vibrations of the stapes footplate, one can find the dynamic amplitude ***A***_***SF***_ of the stapes footplate.

The analysis of the ***inner ear*** is based on two assumptions. The first one is that the power of the sound wave after getting by this wave the round window is absorbed by the vibrations of its elastic membrane. From it yields a direct connection of the amplitude of the round window displacements ***A***_***RW***_ with the amplitude ***A***_***SF***_ of the stapes footplate. The second assumption is that a waving of the basilar membrane has no influence on the amplitude of the sound wave in the cochlea. From it yields that the power of the running wave ***P***_***C***_(***x***) must be equal to the power of the vibrating stapes footplate ***P***_***SF***_. And this in turn enables to state the amplitude of the sound wave in the cochlea ***A***_***C***_(***x***) as a function of the stapes footplate amplitude ***A***_***SF***_. A basal membrane movement depends on the sound wave pressure. Therefore its amplitude ***A***_***BM***_(***y***) is a function of the amplitude of the sound wave ***A***_***C***_(***x***), and finally a function of the amplitude of the stapes footplate ***A***_***SF***_. It is shown that when the sound wave front begins to run in the scala vestibuli, the drop of the sound wave amplitude due to the basilar membrane waving is the order of 10^−6^.

Relations between the wave amplitude, the sound pressure amplitude and the sound wave power are given in the Appendix A. The effect of the basilar membrane waving on the sound wave amplitude is estimated in the Appendix B. The list of the symbols with the basic relations as well as the calculation data one can find in the Appendix C.

## 3. Results

### 3.1. Sound wave in the outer ear

Assume that the sound reaches the ear with a given frequency ***f***_(***i***)_ [1/*s*] and a fixed sound intensity level ***β*** [*dB*]. Then the intensity of sound at the inlet to the ear 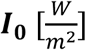 is

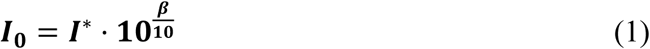

where 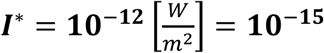 [*N*/(*mm*·*s*)] is the reference sound intensity.

If ***F***_**0**_ [*mm*^2^] is the cross-sectional area of the inlet to the ear canal, then the power of the sound wave ***P***_**0**_ [*N* · *mm*/*s*] which reaches the ear is given by the rule

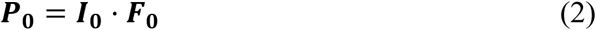

According to our assumptions, ***P***_**0**_ does not change in the whole auditory canal.

Let ***f***_(***i***)_[1/*s*] be a set frequency of the sound wave, 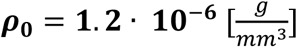 - the air density and 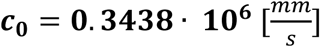 - the sound wave velocity in the air, at 20° C.

Then the amplitude of the wave at the inlet to the ear canal ***A***_**0**_ [*mm*] takes the form (see Eq. (A6))

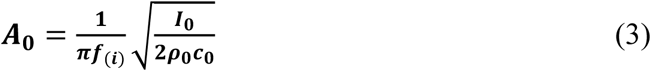

and the amplitude of sound pressure ***p***_**0**_ [*N*/*mm*^2^] corresponding to this wave is (see Eq. (A9)).

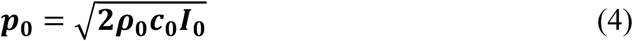

Let us go to the end of the ear canal. Suppose that ***F***_***TM***_ [*mm*^2^] is the cross-sectional area of the ear canal at the base of the tympanic membrane. Thus the sound intensity just before the tympanic membrane ***I***_***TM*−**_ [*N*/(*mm* · *s*)] is.

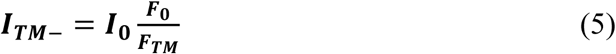

If the value of ***I***_***TM*−**_ is known, one can determine the correlating power of the sound wave and its amplitude. Here, we need only to determine the amplitude of the acoustic pressure ***p***_***TM***_ [*N*/*mm*^2^] acting on the tympanic membrane

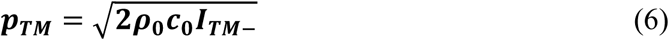

The ***resultant force N***_***TM*−**_ [*N*] of the pressure ***p***_***TM***_ is amplitude of harmonic force acting from the side of the ear canal on the tympanic membrane. Its value is determined by the fixed sound intensity ***I***_**0**_, namely

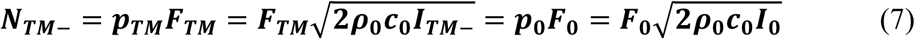

### 3.2. Sound transfer in the middle ear

Here, we will determine the amplitude ***A***_***SF***_ [*mm*] of the stapes footplate vibrations caused by the sound of frequency ***f***_(***i***)_ with the initial power ***P***_**0**_. We will do it in three steps.

1. First, considering the ***reflection*** of part of the sound wave from the tympanic membrane, we will find the amplitude ***N***_***SF***_ of harmonic force acting on the stapes footplate. It will enable us to get the static ***amplitude w***_***SF***_ of the stapes footplate pulsations.
2. Next, an equation of damped forced vibrations, for the stapes footplate which vibrates with a given frequency ***f***_(***i***)_, will be solved. It will enable us to find the ***dynamic amplitude A***_***SF***_ of the stapes footplate vibrations.
3. Finally, we compare the values of ***A***_***SF***_ with the measured amplitudes of the stapes footplate ***d***_***SF***_ got from the temporal bone specimens [8]

#### Ad 1. The static amplitude *w*_*SF*_ of the stapes footplate

The partial reflection of the sound wave from the tympanic membrane implies a damping of the stapes footplate vibrations. This results in the *α*-fold reduction in the sound intensity between the ear canal and the cochlea. The reason for this is a difference between the wave resistance in the air ***Z***_**0**_ **= 0.429·10**^**-6**^ [*N·s/mm*^*3*^] and in the fluid ***Z***_**1**_ **= 1.480·10**^**-3**^ [*N·s/mm*^*3*^]. The quantity ***α*** is the absorption coefficient of the sound wave power

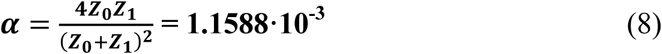

and it has the greatest impact on the attenuation of sound in the middle and inner ear. Due to the partial reflection of the sound wave, the sound intensity ***I***_***TM*−**_ drops in the middle ear to the value

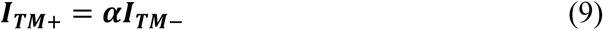

Thus, the maximum force ***N***_***TM***+_ [*N*] with which the tympanic membrane acts on the ossicles is

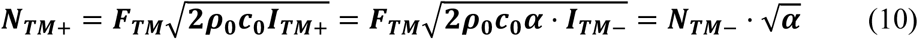

The malleus-incus pair creates a lever increasing the ***γ*** = **1. 3**-fold force acting on the stapes footplate. Denoting this force by ***N***_***SF***_ [*N*], one can obtain

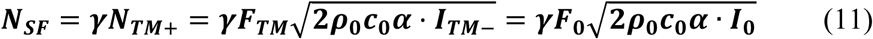

Assume for a moment the quasi-static pulsations of the stapes footplate. In this case, the viscosity of the annular ligament has no influence on the stapes footplate amplitude ***w***_***SF***_. The amplitude ***w***_***SF***_ depends only on the force ***N***_***SF***_ and the stiffness of the annular ligament 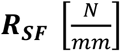. According to [9], we assume 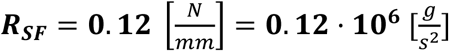. So, the ***static amplitude w***_***SF***_ has the form

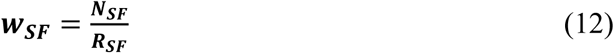

Note that the quasi-static amplitude ***w***_***SF***_ does not depend on the frequency ***f***_(***i***)_ of the force with the amplitude ***N***_***SF***_, which moves the stapes footplate. In the Appendix C, one can find typical dimensions of the external and middle ear. For these dimensions and a level of the sound intensity from the range **10 *dB*** ≤ ***β*** ≤ **120 *dB***, the values of the quasi-static amplitude ***w***_***SF***_ are shown in Table1.

**Table 1.**
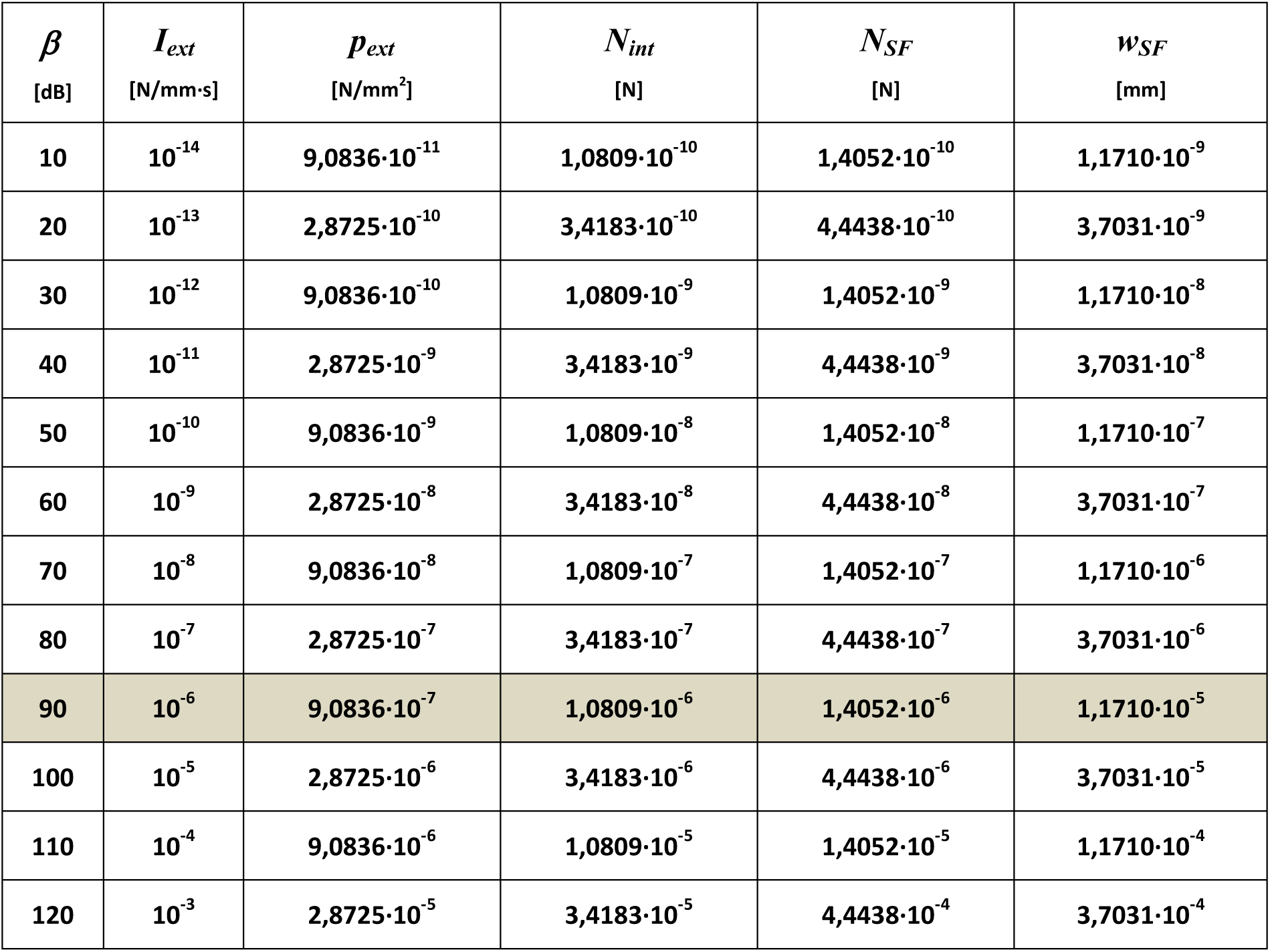

| $\beta$<br>[dB] | $I_{ext}$<br>[N/mm·s] | $p_{ext}$<br>[N/mm<sup>2</sup>] | $N_{int}$<br>[N] | $N_{SF}$<br>[N] | $w_{SF}$<br>[mm] |
| --- | --- | --- | --- | --- | --- |
| 10 | $10^{-14}$ | $9,0836 \cdot 10^{-11}$ | $1,0809 \cdot 10^{-10}$ | $1,4052 \cdot 10^{-10}$ | $1,1710 \cdot 10^{-9}$ |
| 20 | $10^{-13}$ | $2,8725 \cdot 10^{-10}$ | $3,4183 \cdot 10^{-10}$ | $4,4438 \cdot 10^{-10}$ | $3,7031 \cdot 10^{-9}$ |
| 30 | $10^{-12}$ | $9,0836 \cdot 10^{-10}$ | $1,0809 \cdot 10^{-9}$ | $1,4052 \cdot 10^{-9}$ | $1,1710 \cdot 10^{-8}$ |
| 40 | $10^{-11}$ | $2,8725 \cdot 10^{-9}$ | $3,4183 \cdot 10^{-9}$ | $4,4438 \cdot 10^{-9}$ | $3,7031 \cdot 10^{-8}$ |
| 50 | $10^{-10}$ | $9,0836 \cdot 10^{-9}$ | $1,0809 \cdot 10^{-8}$ | $1,4052 \cdot 10^{-8}$ | $1,1710 \cdot 10^{-7}$ |
| 60 | $10^{-9}$ | $2,8725 \cdot 10^{-8}$ | $3,4183 \cdot 10^{-8}$ | $4,4438 \cdot 10^{-8}$ | $3,7031 \cdot 10^{-7}$ |
| 70 | $10^{-8}$ | $9,0836 \cdot 10^{-8}$ | $1,0809 \cdot 10^{-7}$ | $1,4052 \cdot 10^{-7}$ | $1,1710 \cdot 10^{-6}$ |
| 80 | $10^{-7}$ | $2,8725 \cdot 10^{-7}$ | $3,4183 \cdot 10^{-7}$ | $4,4438 \cdot 10^{-7}$ | $3,7031 \cdot 10^{-6}$ |
| 90 | $10^{-6}$ | $9,0836 \cdot 10^{-7}$ | $1,0809 \cdot 10^{-6}$ | $1,4052 \cdot 10^{-6}$ | $1,1710 \cdot 10^{-5}$ |
| 100 | $10^{-5}$ | $2,8725 \cdot 10^{-6}$ | $3,4183 \cdot 10^{-6}$ | $4,4438 \cdot 10^{-6}$ | $3,7031 \cdot 10^{-5}$ |
| 110 | $10^{-4}$ | $9,0836 \cdot 10^{-6}$ | $1,0809 \cdot 10^{-5}$ | $1,4052 \cdot 10^{-5}$ | $1,1710 \cdot 10^{-4}$ |
| 120 | $10^{-3}$ | $2,8725 \cdot 10^{-5}$ | $3,4183 \cdot 10^{-5}$ | $4,4438 \cdot 10^{-4}$ | $3,7031 \cdot 10^{-4}$ |

#### Ad 2. The dynamic amplitude *A*_*SF*_ of the stapes footplate

For a fixed level of the sound intensity ***β* = 90 *dB***, one can get that ***w***_***SF***_ = **11. 7 *nm*** = **1. 17 10**^−**8**^ ***m***. The obtained value is close to that obtained from measurements of the stapes footplate amplitude for ***β* = 90 *dB*** and the sound frequency ***f***_(***i***)_ = **1000 *Hz*** [8]. It is the value of ***d***_***SF***_ = **10. 2 *nm*** = **10. 2 10**^−**8**^ ***m***. One can show that ***f***_(***i***)_ = **1000 *Hz*** is the ***natural frequency*** of the stapes footplate, so in the absence of damping, a resonance should take place. In the case of damping, for ***k***_***SF***_ = **2*πf***_(***i***)_, the dynamic amplitude of vibrations ***A***_***SF***_ is the largest one and close to the static amplitude ***w***_***SF***_. Then, one can take an ***additional assumption***:

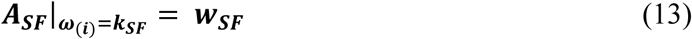

Denote by ***M***_***SF***_ the mass of the stapes suspended on the annular ligament. Recall, that the stapes footplate is subjected to the vibrations forced by a harmonic force ***N***(***t***) = ***N***_***SF***_ **sin**(***ω***_(***i***)_***t***) and damped with a parameter ***D***_***SF***_. Here, ***ω***_(***i***)_ = **2*πf***_(***i***)_) [1*/s*] is the circular frequency of the tested vibration frequency ***f***_(***i***)_. The equation of vibration of the stapes footplate has the form

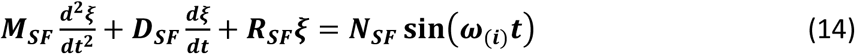

where ***ξ = ξ* (*t*)** [*mm*] - the position of the front of the footplate with respect to the equilibrium position. It may be written as

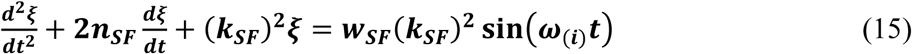

where 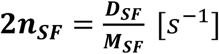 is a damping factor and 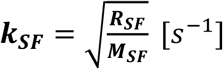 – the ***resonant angular frequency*** of the stapes footplate. A solution of the last equation gives rules for the footplate displacements

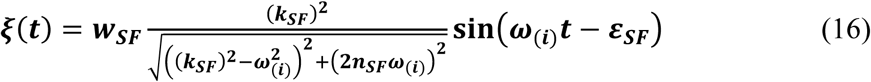

for

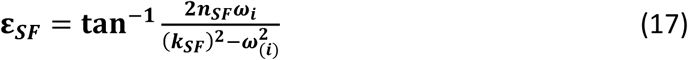

Thus, the following expression for the ***dynamic amplitude*** of the stapes footplate is given by the fixed sound frequency

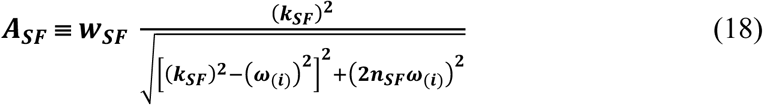

Notice that 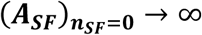, when ***ω***_***i***_ = ***k***_***SF***_ =**6.283·10**^**3**^ ***s***^***-1***^ and 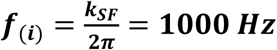, it take place:

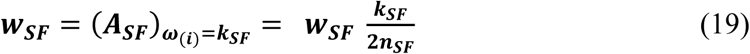

Thus, the damping factor **2*n***_***SF***_ is equal to:

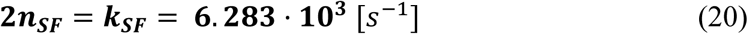

And because 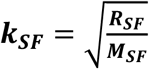, the corresponding mass of the stapes is equal to **3. 04 · 10**^−**3**^ [g]. This value is known as the average measured mass of the stapes.

Now, the expression for the dynamic amplitude of the stapes footplate takes the form:

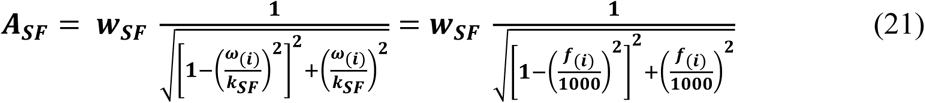

The effect of dynamics on the amplitude of the stapes shield shows the function **20 log**_**10**_((***A***_***SF***_(***f***_(***i***)_)**/**(***w***_***SF***_), which is given for **0** ≤ ***f***_(***i***)_ ≤ **20000** *Hz* at Fig. 3.

**Fig. 1.**
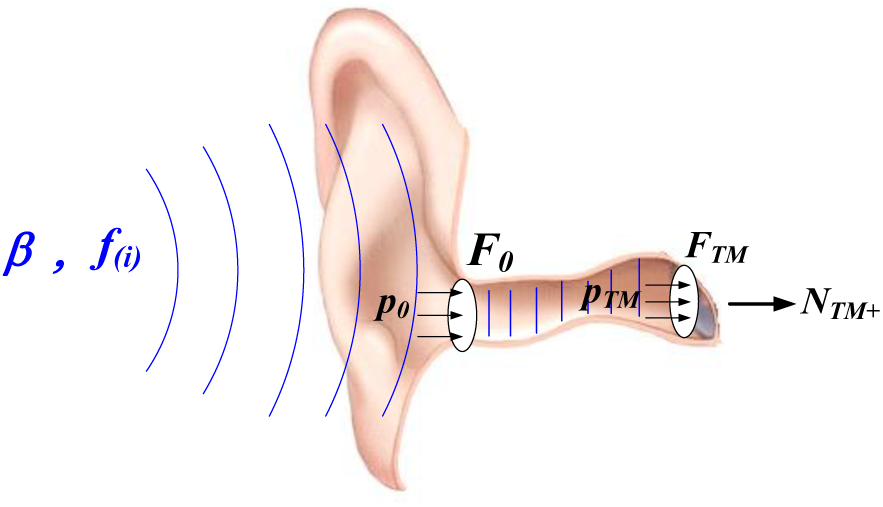
Sound wave propagation in the outer ear.

**Fig. 2.**
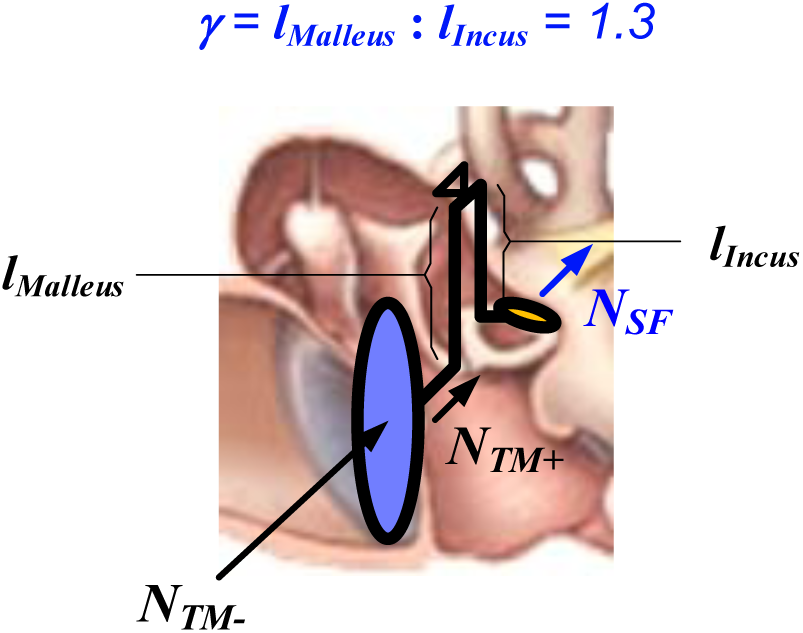
Sound transfer in the middle ear.

**Fig. 3.**
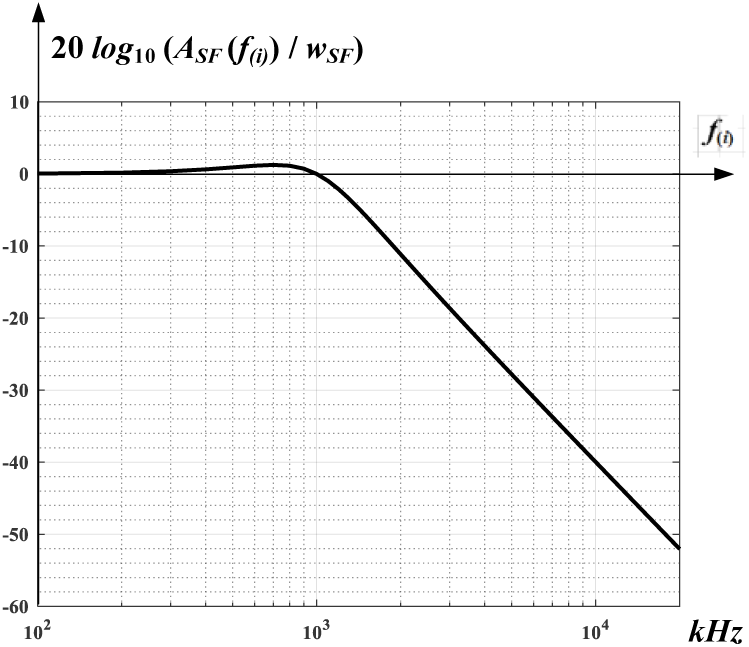
Function ln(*A*_*SF*_/*w*_*SF*_) vs sound frequency *f*_(*i*_).

#### Comparison of the amplitude *A*_*SF*_ with the measurements results *d*_*SF*_

Values of the measured amplitudes of the stapes footplate are given in [8]. The tests were done with help of human temporal bones by exposing the tympanic membrane to the fixed sound intensity level of **90** *dB*. The amplitudes of the stapes footplate ***d***_***SF***_ were measured for a set of frequency from the range **400-10000** Hz. The results are shown in Table 2.

**Table 2.**

| Test<br>*i* | Frequency<br>$f(i)$ [Hz] | Measured amplitude<br>$d_{SF}$ [mm] | Calculated amplitude<br>$A_{SF}$ [mm] | Relative difference<br>$\frac{A_{SF}-d_{SF}}{d_{SF}} \cdot 100$ [%] |
| --- | --- | --- | --- | --- |
| 1 | 400 | $1.64 \cdot 10^{-5}$ | $1.2576 \cdot 10^{-5}$ | 23 |
| 2 | 500 | $1.53 \cdot 10^{-5}$ | $1.2980 \cdot 10^{-5}$ | 15 |
| 3 | 630 | $1.90 \cdot 10^{-5}$ | $1.3415 \cdot 10^{-5}$ | 29 |
| 4 | 800 | $1.97 \cdot 10^{-5}$ | $1.3337 \cdot 10^{-5}$ | 32 |
| 5 | 1000 | $1.02 \cdot 10^{-5}$ | $1.1700 \cdot 10^{-5}$ | 15 |
| 6 | 1250 | $8.92 \cdot 10^{-6}$ | $8.5351 \cdot 10^{-6}$ | 4 |
| 7 | 1600 | $7.12 \cdot 10^{-6}$ | $5.2354 \cdot 10^{-6}$ | 27 |
| 8 | 2000 | $4.93 \cdot 10^{-6}$ | $3.2448 \cdot 10^{-6}$ | 34 |
| 9 | 2500 | $2.84 \cdot 10^{-6}$ | $2.0120 \cdot 10^{-6}$ | 29 |
| 10 | 3150 | $6.40 \cdot 10^{-7}$ | $1.2364 \cdot 10^{-6}$ | 93 |
| 11 | 4000 | $7.73 \cdot 10^{-7}$ | $7.5362 \cdot 10^{-7}$ | 2 |
| 12 | 5000 | $8.36 \cdot 10^{-7}$ | $4.7722 \cdot 10^{-7}$ | 43 |
| 13 | 6300 | $6.14 \cdot 10^{-7}$ | $2.9845 \cdot 10^{-7}$ | 51 |
| 14 | 8000 | $1.69 \cdot 10^{-7}$ | $1.8422 \cdot 10^{-7}$ | 9 |
| 15 | 10000 | $7.57 \cdot 10^{-8}$ | $1.1757 \cdot 10^{-7}$ | 55 |

Note that the differences between calculated and measured values vary from **2%** to **93%**. A mean difference does not exceed **31%**. Taking into account the difficulty in taking measurements, the differences seem to be acceptable. Due to the more low accuracy of the measured values, these differences seem to be acceptable.

### 3.3. Round window membrane behavior as a response to the stapes footplate vibrations

A travel of the sound wave ends at the ***elastic membrane*** of the round window. The round window membrane vibrates due to the pulsation of the sound wave pressure (see Fig. 4). Now, dynamic amplitude of the vibrations of the round window is looked for. To do it, one need to determine the membrane parameters and use of the assumption that it absorbs all the energy of the sound wave.

**Fig. 4.**
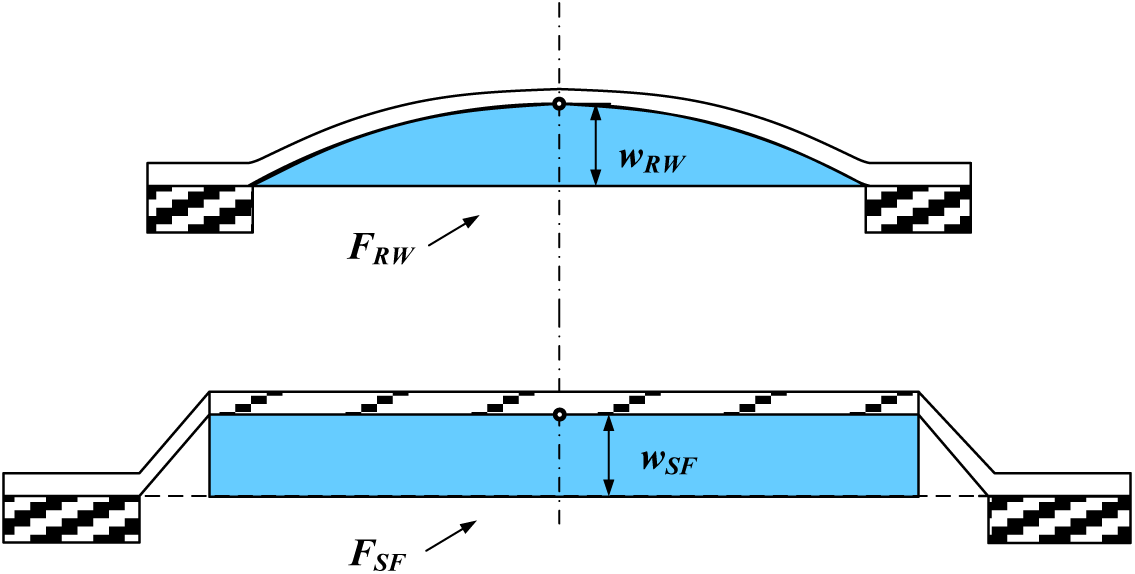
Motion of the round window membrane.

Let ***r***_***RW***_ be the radius of the round window membrane, ***R***_***RW***_ **−** its stiffness and ***A***_***RW***_ **−** the amplitude of its vibrations. To find the stiffness ***R***_***RW***_ of the membrane, the following reasoning is done. Let us assume that the deflected round window membrane has the shape of paraboloid with static amplitude ***w***_***RW***_. Note that the volume of fluid contained in it is half the volume of the cylinder with the height ***w***_***RW***_.

Assume that a quasi-static force ***N***_***SF***_ moves the stapes footplate on a distance ***w***_***SF***_. Then the fluid in the cochlea will push the center of the round window membrane, with the same force, on a distance ***w***_***RW***_ (see Fig.4).

Comparing the volume of the fluid pushed through the stapes footplate with the volume of the fluid contained in the deflected membrane, one can get a relation between ***w***_***RW***_ and ***w***_***SF***_, namely

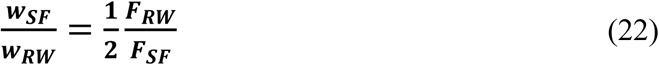

At every point inside the cochlea, the static pressure caused by the given force ***N***_***SF***_ is the same and equal to ***p***_***s***_ = ***N***_***SF***_**/*F***_***SF***_. Thus, on the round window membrane acts the force

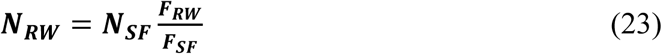

Because ***N***_***SF***_ = ***R***_***SF***_ ***w***_***SF***_ and ***N***_***RW***_ = ***R***_***RW***_ ***w***_***RW***_, one can obtain

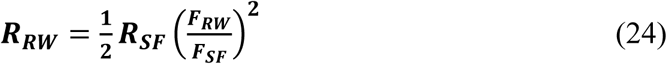

For parameters ***F***_***SF***_ and ***F***_***RW***_ given in the Appendix C, we have: 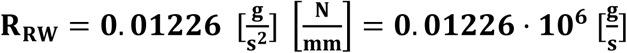.

Recall that in each cross-section of the cochlea, the total power transmitted by the stapes footplate is the same and equal to ***P***_***SF***_. Thus, the power of the round window membrane ***P***_***RW***_ is equal to this quantity

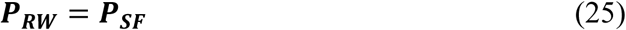

- **The power of the stapes footplate *P***_***SF***_ The power ***P***_***SF***_ of the stapes footplate can be defined as the work done by during the time ***T***_(***i***)_ of one vibration cycle. The work done by the footplate in time 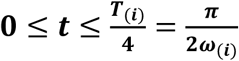, during ¼ of the vibration cycle is

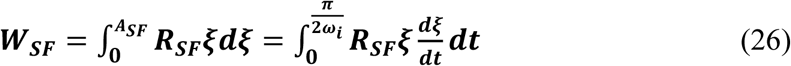

and the ***power of the stapes footplate*** is the work **2*W***_***SF***_ done in the time ***T***_(***i***)_. Then, we have:

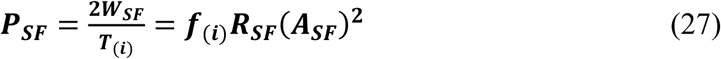
- **The power of the round window membrane *P***_***RW***_ The power of the membrane ***P***_***RW***_ is calculated in the same way as the power of the vibrating stapes footplate, namely

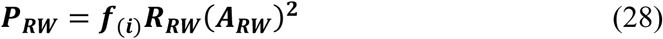

Comparing Eq. (27) and (28) one can get

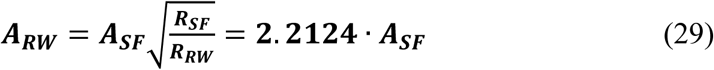

Values of the measured amplitudes of the round window membrane are given in [9]. The tests were done with help of human temporal bones by exposing the tympanic membrane to the fixed sound intensity level of **90** *dB*. The amplitudes of the stapes footplate ***d***_***RW***_ were measured for a set of four frequencies from the range **1000-8000** Hz. The results are shown in Table 3.

**Table 3.**

| No | Frequency<br>$f_i$<br>[Hz] | Amplitude of<br>stapes<br>footplate $A_{SF}$<br>[mm] | Amplitude of<br>round window<br>membrane $A_{RW}$<br>[mm] | Measured<br>dynamic RW<br>amplitude | Relative<br>difference<br>[%] |
| --- | --- | --- | --- | --- | --- |
| 1 | 1000 | $1.1700 \cdot 10^{-05}$ | $2.5886 \cdot 10^{-05}$ | $2.5 \cdot 10^{-05}$ | 4 |
| 2 | 2000 | $3.2448 \cdot 10^{-06}$ | $0.7180 \cdot 10^{-05}$ | $1.0 \cdot 10^{-05}$ | 28 |
| 3 | 4000 | $7.5362 \cdot 10^{-07}$ | $1.6674 \cdot 10^{-06}$ | $1.6 \cdot 10^{-06}$ | 4 |
| 4 | 8000 | $1.8422 \cdot 10^{-07}$ | $0.4076 \cdot 10^{-06}$ | $0.6 \cdot 10^{-06}$ | 32 |

### 3.3. Sound wave travel in the inner ear

Geometry of the assumed model of the inner ear is shown at Fig. 5. The dimensions of the cochlea which appear in the derived rules are marked there. Their values used for the calculations are given in Appendix C.

**Fig. 5.**
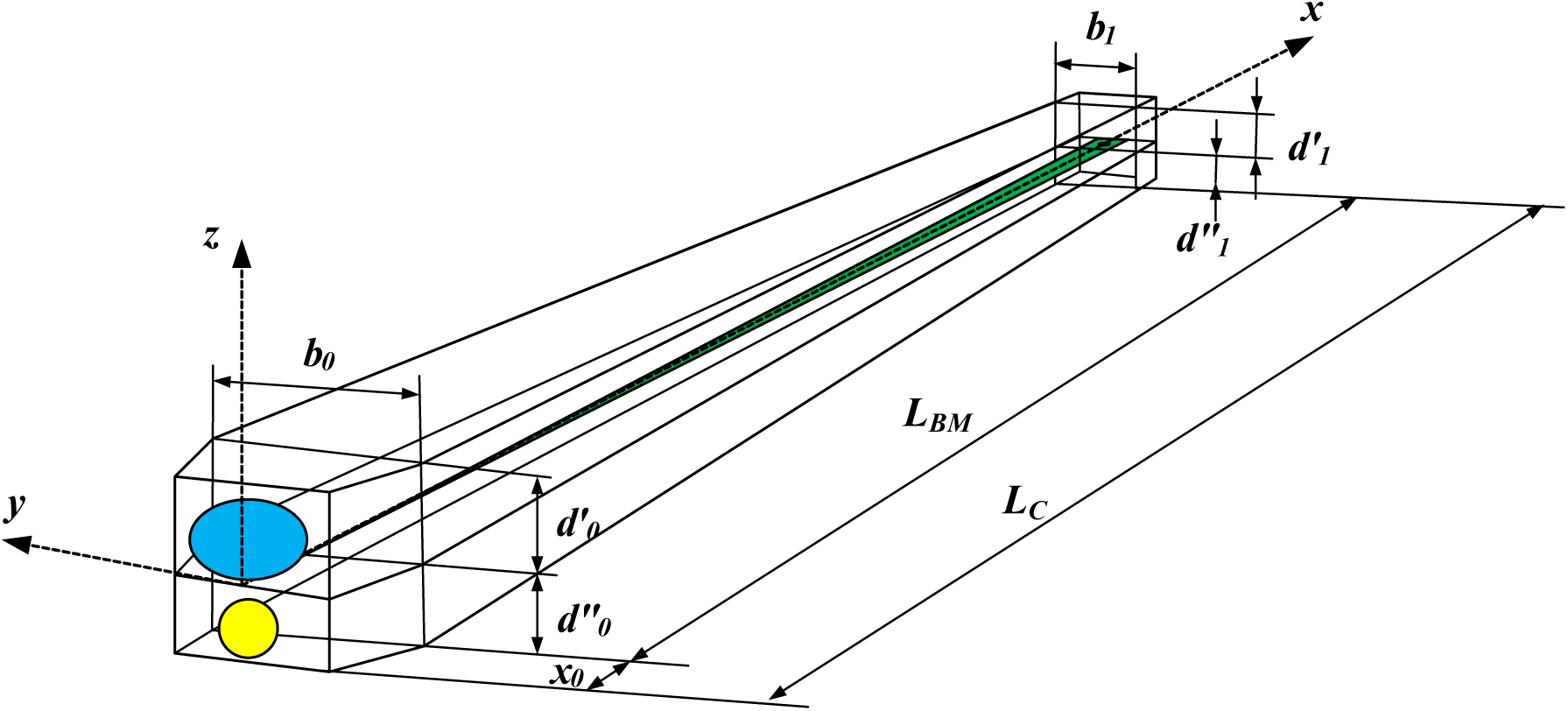
Geometry of the inner ear.

Recall that the inner ear is a waveguide of the length **2*L***_***C***_ with a variable cross-section ***F***_***C***_(***x***) filled with fluid, with the stapes footplate at the inlet and with the round window membrane at the end. The ***x***-axis of the waveguide is straight, but at the middle of the waveguide it changes direction to the opposite one. In the fluid runs an ***elastic plane wave*** forced by the vibrations of the stapes footplate with the amplitude ***A***_***SF***_. After reaching the membrane of the round window, the sound wave is completely suppressed by it. In place of the basilar membrane we take a chain of uncoupled viscoelastic fibers that move under the pressure of the running wave. The ***y***-axis of the fiber chain up to half of the double channel waveguide overlaps the ***x***-axis. It begins at the base of the fiber chain, at a distance ***x***_**0**_ from the oval window and ends with a hole just before the cochlea apex. The width ***a***(***y***) and the thickness ***h***(***y***) of the basilar membrane as well as its average density ***ρ***_***BM***_ and viscosity ***d***_***BM***_(***y***) are known.

Let ***A***_***C***_(***x***) is amplitude of the sound wave and ***A***_***BM***_(***x***) **−** amplitude of the chain of the fibers. The expression of these values by the frequency ***f***_(***i***)_ and the intensity level ***β*** of the sound wave reaching the ear will be done in two steps.

#### 1. Dependence of the sound wave amplitude on the stapes footplate amplitude

Because the basilar membrane motion has no influence on the amplitude of the sound wave, one can state that

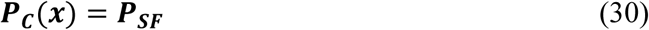

- **The power of the running wave *P***_***C***_(***x***) Consider a wave front which runs in the cochlea with an amplitude ***A***_***C***_(***x***) and frequency ***f***_(***i***)_. A position of the wave front at time ***t*** is ***x*** = ***x***(***t***) = ***c***_**1**_***t***, with ***c***_**1**_ as the wave velocity in the fluid of density ***ρ***_**1**_. The cross-section of the waveguide on the wave front is ***F***_***C***_(***x***). Using the Eq. (A6), one can formulate the expression for the ***wave power P***_***C***_(***x***) as a function of the distance ***x*** from the oval window

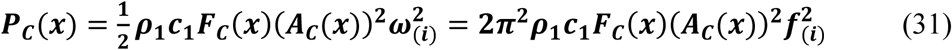

The above expression is valid for **0** ≤ ***x*** ≤ **2*L***_***C***_, and so both in the scala vestibuli and in the scala tympani. Comparing Eqs (31) and (27), one can show the sound wave amplitude ***A***_***C***_(***x***) as a function of the stapes footplate amplitude ***A***_***SF***_, namely

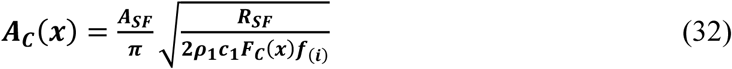

Notice that each point ***y*** of the basilar membrane has a coordinate ***x***_**0**_ ≤ ***x***′ ≤ ***x***_**0**_ + ***L***_***BM***_ when viewed from scala vestibuli and the coordinate ***x***_**0**_ + ***L***_***BM***_ ≤ ***x*″** ≤ ***x***_**0**_ + **2*L***_***BM***_ when viewed from scala tympani. The corresponding cross-sections of the scala vestibuli and the scala tympani are ***F***_***C***_(***x*′**) and ***F***_***C***_(***x*″**), respectively. Thus

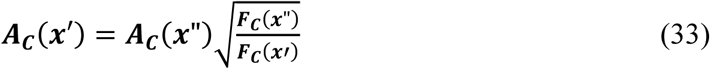

At the oval window the sound wave amplitude is 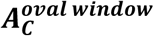 is equal 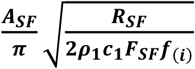. One can see that 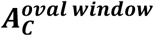 is much smaller than ***A***_***SF***_. Going forward along the scala vestibuli, at the basilar membrane base, one can find the sound wave amplitude 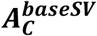. Passing through the cochlea apex to the scala tympani, one can observe a step change in the cross-section, which results in a slight increase in the amplitude of the sound wave equal to 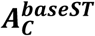. In fact, taking into account the loss of energy of the wave due to the movement resistances, the amplitude increase should be smaller.

Using the cochlea dimensions taken in the Appendix C, one can show differences between 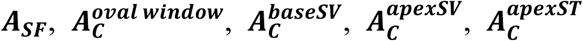 and 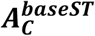. In the Table 4 the suitable values are shown for a set of four frequencies from the range **1000-8000** Hz.

**Table 4.**

| $N_o$ | $f_{(i)}$<br>[Hz] | $A_{SF}$<br>[mm] | $A_C^{oval\ window}$<br>[mm] | $A_C^{baseSV}$<br>[mm] | $A_C^{apexSV}$<br>[mm] | $A_C^{apexST}$<br>[mm] | $A_C^{baseST}$<br>[mm] |
| --- | --- | --- | --- | --- | --- | --- | --- |
| 1 | 1000 | $1.1700 \cdot 10^{-05}$ | $4.7749 \cdot 10^{-07}$ | $2.6958 \cdot 10^{-07}$ | $6.4577 \cdot 10^{-07}$ | $7.7680 \cdot 10^{-07}$ | $3.1173 \cdot 10^{-07}$ |
| 2 | 2000 | $3.2448 \cdot 10^{-06}$ | $9.3638 \cdot 10^{-08}$ | $5.2866 \cdot 10^{-08}$ | $1.2664 \cdot 10^{-07}$ | $1.5233 \cdot 10^{-07}$ | $6.1131 \cdot 10^{-08}$ |
| 3 | 4000 | $7.5362 \cdot 10^{-07}$ | $1.5378 \cdot 10^{-08}$ | $8.6822 \cdot 10^{-09}$ | $2.0797 \cdot 10^{-08}$ | $2.5017 \cdot 10^{-08}$ | $1.0040 \cdot 10^{-08}$ |
| 4 | 10000 | $1.1757 \cdot 10^{-07}$ | $1.5173 \cdot 10^{-09}$ | $8.5665 \cdot 10^{-10}$ | $2.0521 \cdot 10^{-09}$ | $2.4684 \cdot 10^{-09}$ | $9.9058 \cdot 10^{-10}$ |

#### 2. Dependence of the basilar membrane amplitude on the stapes footplate amplitude

The chain of fibers with the length of ***L***_***BM***_, taken as the basilar membrane, runs along the ***y*** axis, which partially overlaps the ***x***-axis. Its base lies at a distance ***x***_**0**_ from the oval window and ends before the cochlea apex. Thus, the relationship between the ***x*** and ***y*** axes is as follows

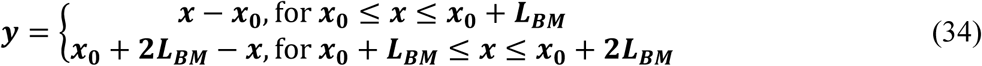

Let us consider a point ***y*** of the basilar membrane. This point has a coordinate ***x*′** in the scala vestibuli and a coordinate ***x*″** in the scala tympani. One can see that ***x*″** = ***x*′** + **Δ*X***, where

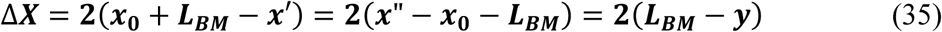

The sound wave acts on the basilar membrane at the point y due to a difference between the sound wave pressures at this point in the vestibuli scala and scala tympani. Let us consider a point ***y*** of the basilar membrane with a coordinate ***x*′** in the ***scala vestibuli*** and a coordinate ***x*″** in the ***scala tympani***. The wave front passes through the coordinate ***x*′** in the scala vestibuli at time ***t***. A deflection of the fluid particle over the coordinate ***x*′** is equal to the amplitude ***ξ***_***C***_(***x***′, ***t***) = ***A***_***C***_(***x*′**). After the time **Δ*t*** = **Δ*X***(***x***′)/***c***_**1**_ the wave front reaches the coordinate ***x*″** in the scala tympani. Now the deflection of the particle below the coordinate ***x*″** is equal to the amplitude ***ξ***_***C***_(***x*″, *t*** + **Δ*t***) = ***A***_***C***_(***x*″**). In the meantime the deflection of the particle over the coordinate ***x*′** will drop to the value ***ξ***_***C***_(***x*′, *t*** + **Δ*t***) = ***A***_***C***_(***x*′**) ***ρ***_**1**_***c***_**1**_***ω***_(***i***)_ **sin**(***ω***_***i***_**Δ*X***(***x***′)/***c***_**1**_) (see Eq. (A1)).

Thus, the sound pressures at the point ***y*** and at the time ***t*** + **Δ*t*** is equal to (see Eq. (A8))

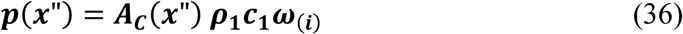

when the wave front runs through the scala tympani, and

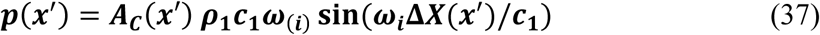

for the wave running in the scala vestibuli. Here ***ρ***_**1**_ is the cochlea fluid density, ***c***_**1**_ is the speed of the sound wave, ***ω***_(***i***)_ is the circular sound frequency and

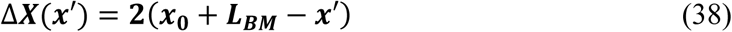

Concluding, one can state that at each point **0** ≤ ***y*** ≤ ***L***_***BM***_ of the basilar membrane acts a ***differential pressure***

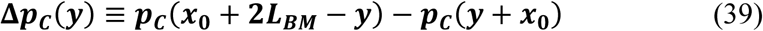

Takin into account Eq. (33), one can introduce a ***differential amplitude*** at the point ***y***

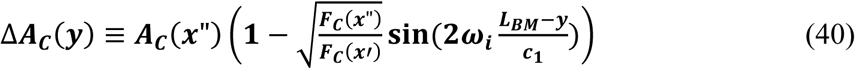

for

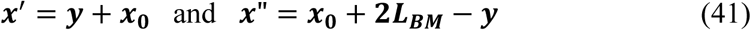

Because the sound wave amplitude depends on the stapes footplate amplitude as it is stated by Eq. (32), Eq. (40) takes the form

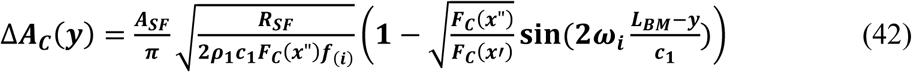

Thus, Eq. (39) takes the form

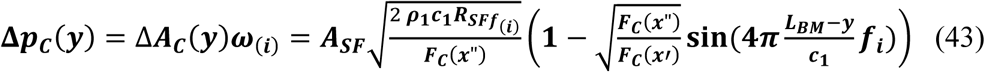

Now, consider a single fiber of width **Δ*y*** cut out from the membrane, fastened at both ends and loaded on the surface ***a***(***y***) **· Δ*y*** by the sound wave differential pressure **Δ*p***_***C***_(***y***). Resultant force acting on the fiber with unite width is equal to ***N***_***BM***_(***y***) = **Δ*p***_***C***_(***y***) ***a***(***y***) **Δ*y***. Because the properties of the fiber are known, one can find its static deflection ***u***_**0**_(***y***), and next the normalized with respect to **Δ*y***, rigidity of the fiber ***r***_***BM***_(***y***) = **Δ*p***_***C***_(***y***) ***a***(***y***) /***u***_**0**_(***y***). Notice that, ***normalized mass*** of the fiber is ***m***_***BM***_(***y***) = ***ρ***_***BM***_***a***(***y***)***h***(***y***), and its ***normalized damping factor*** is equal to the viscosity of the basilar membrane ***d***_***BM***_(***y***).

The equation of the vibrations of the mass ***m***_***BM***_(***y***) suspended in the middle of the fiber is in the form

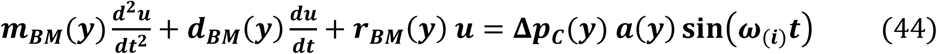

where ***u*** = ***u***(***y, t***) is a dynamic displacement of the mass ***m***_***BM***_(***y***).

Having Eq. (42) one can write

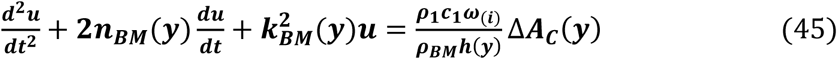

Here 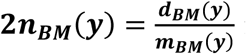 is damping coefficient of the fiber at the point ***y*** for **0** ≤ ***y*** ≤ ***L***_***BM***_, and 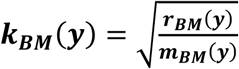 is its ***resonant angular frequency***.

A distribution of the resonant frequencies of the fibers of the basilar membrane along its length is shown by the function ***f***_***BM***_(***y***) **= *k***_***BM***_(***y***)**/**(**2*π***).

Notice that the function ***f***_***BM***_(***y***) can be found if the dimensions and the mechanical properties of the basilar membrane are known. It is also possible to use the form of ***f***_***BM***_(***y***) given in the literature [16]. Here, this second option will be used.

The general form of the solution sought for equation (45) is as follows.

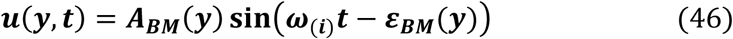

where ***A***_***BM***_(***y***) is an unknown amplitude of the basilar membrane, and ***ε***_***BM***_(***y***) is an unknown phase shift. The real solution of the equation (44) is

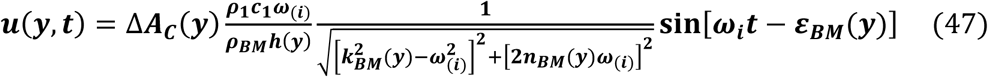

for

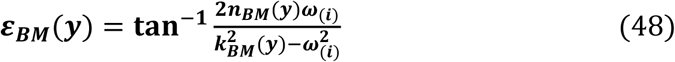

From Eqs (46) and (47) yields the relation between the amplitude of the basal membrane ***A***_***BM***_(***y***) and the differential amplitude of the sound wave **Δ*A***_***C***_(***y***).

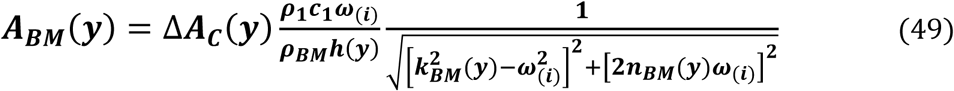

Using Eqs (42) in Eq. (49), a dependence of the basilar membrane on the stapes footplate amplitude takes the form

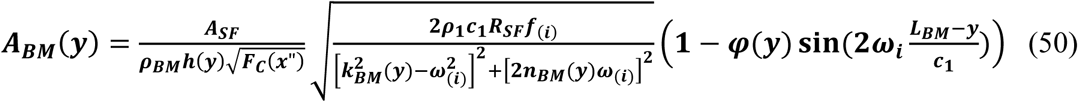

where

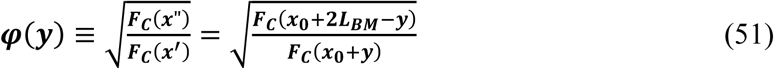

### 3.4. Basilar membrane behavior

#### Boundary conditions

The displacements of the basilar membrane ***u***(***y, t***) should meet the conditions [14]:

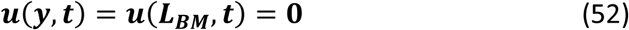

From Eq. (51) yields that the conditions (52) will be satisfied if instead of the thickness of the basilar membrane ***h***(***y***), it will be assumed the function ***h***(***y***)***χ***(***y***)***χ***(***L***_***C***_ − ***y***), where

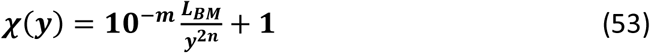

Here, it is taken *m* = **3** and ***n*** = 1. The new thickness function tends to infinity in a surrounding of the basilar membrane ends. So, the displacements ***u***(***y, t***) tend to zero at these points.

#### Resonant frequencies distribution

As resonance frequencies 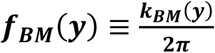 one can take the function proposed by Greenwood in [16]

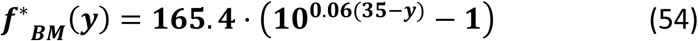

or by Kwacz et al. [8]

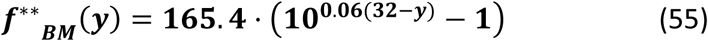

for **0** ≤ ***y*** ≤ ***L***_***BM***_, measured in millimeters. Here, the following function is taken (Fig.7)

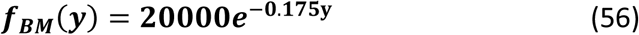

**Fig. 6.**
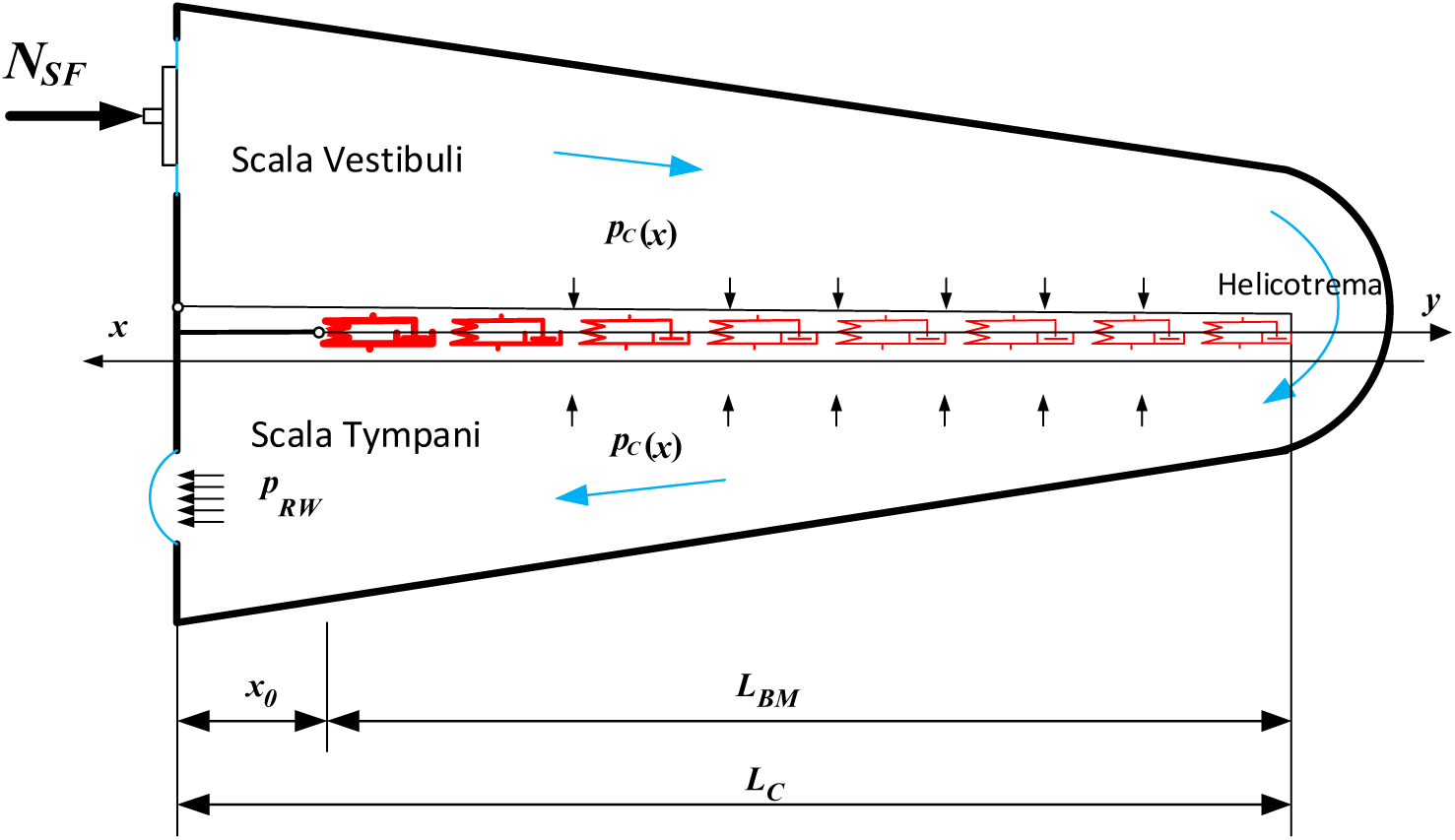
Sound wave propagation in the inner ear.

**Fig. 7.**
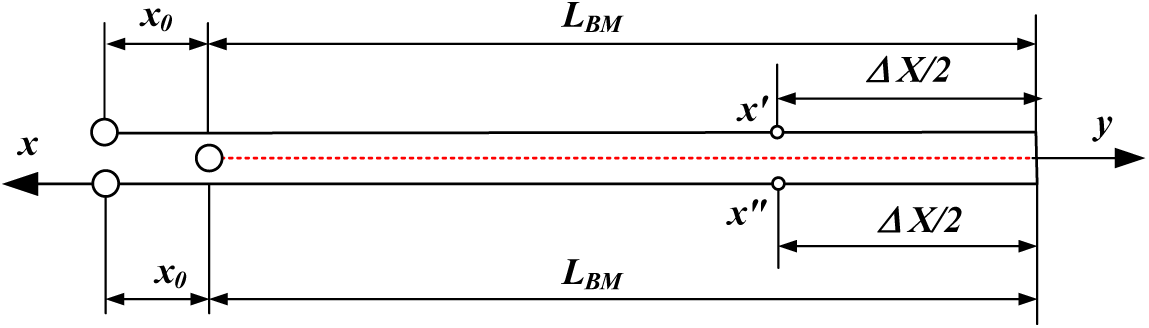
Assumed model of basilar membrane.

#### Level of cochlear amplification

Consider a fixed point ***y***^∗^ of the basilar membrane with the coordinate in the tympanic duct ***x***^∗^ = **2*L***_***BM***_ + ***x***_**0**_ − ***y***^∗^ (see Eq. (41b)). The front of the sound wave reaches the point ***y***^∗^ at the time 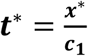. Thus, the amplitude of the basilar membrane displacement at this point (Eq. (51)) may be shown as a function of the sound wave frequency ***f***_(***i***)_ and the fixed parameter ***y***^∗^

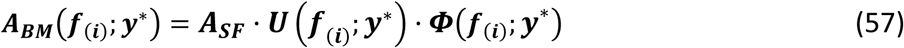

where

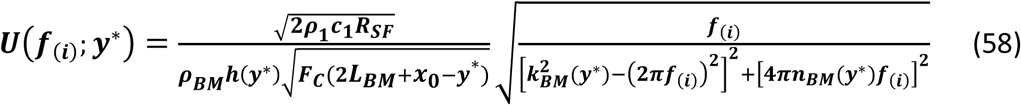

and

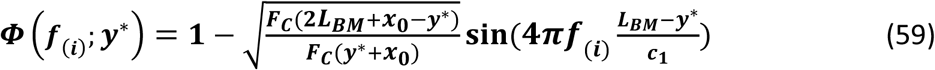

Let us consider the ratio of the amplitude of the basilar membrane ***A***_***BM***_(***f***_(***i***)_**; *y***^∗^) at the fixed point ***y***^∗^ to the amplitude of the stapes footplate ***A***_***SF***_(***f***(***i***)) for the time 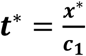. The function

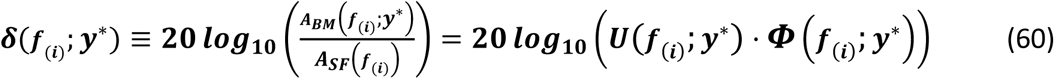

shows a ***level of cochlear amplification*** at the point ***y***^∗^.

The results of measurements of the velocity of the basilar membrane in the tympanic duct are shown in the work [15]. The tests were done at the point located at ***y***^∗^ = **12** [*mm*] from the oval window for different sound wave frequencies. The results were given in ***dB*** and normalized with the velocity of the stapes footplate. To compare them with those given by the rule (60), one can use the data from Appendix B (see Fig. 9).

**Fig. 8.**
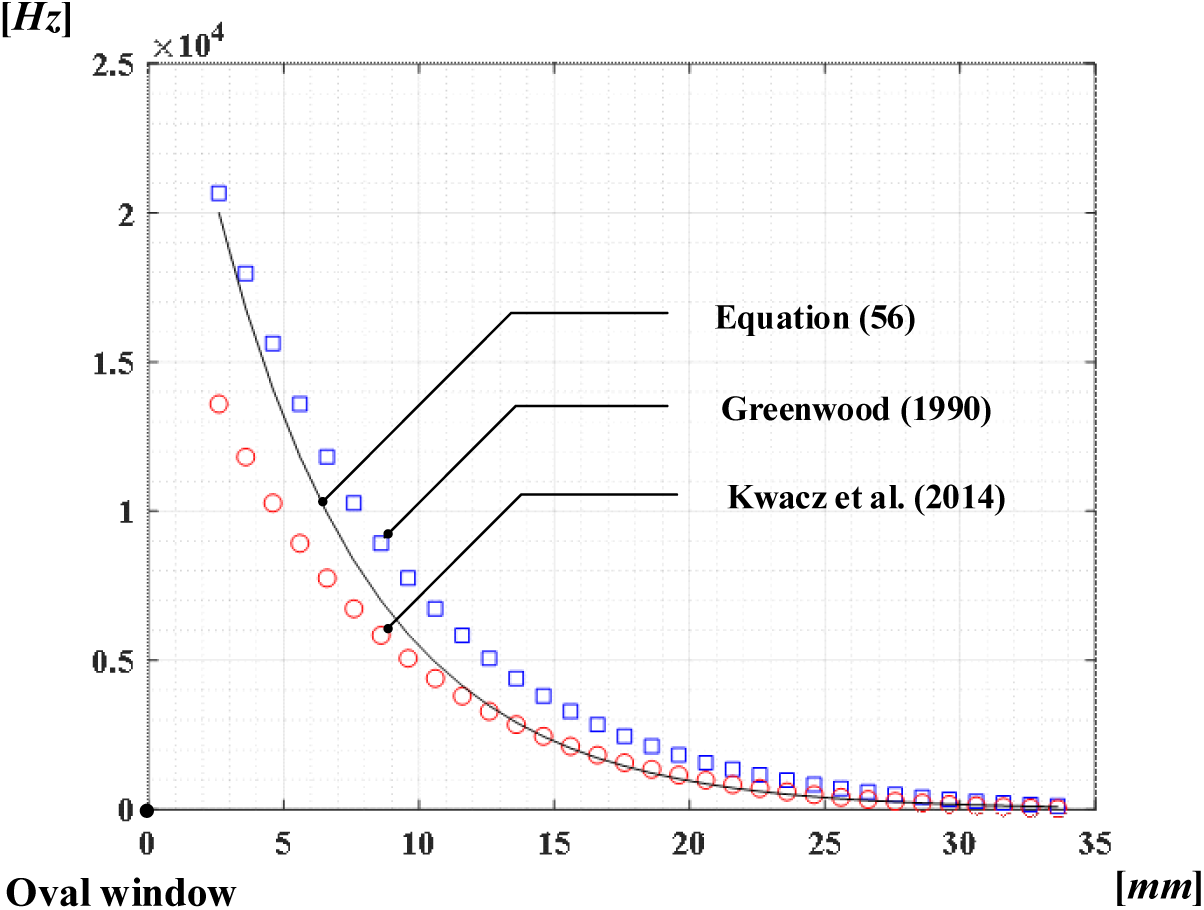
Resonant frequencies of the basilar membrane.

**Fig. 9.**
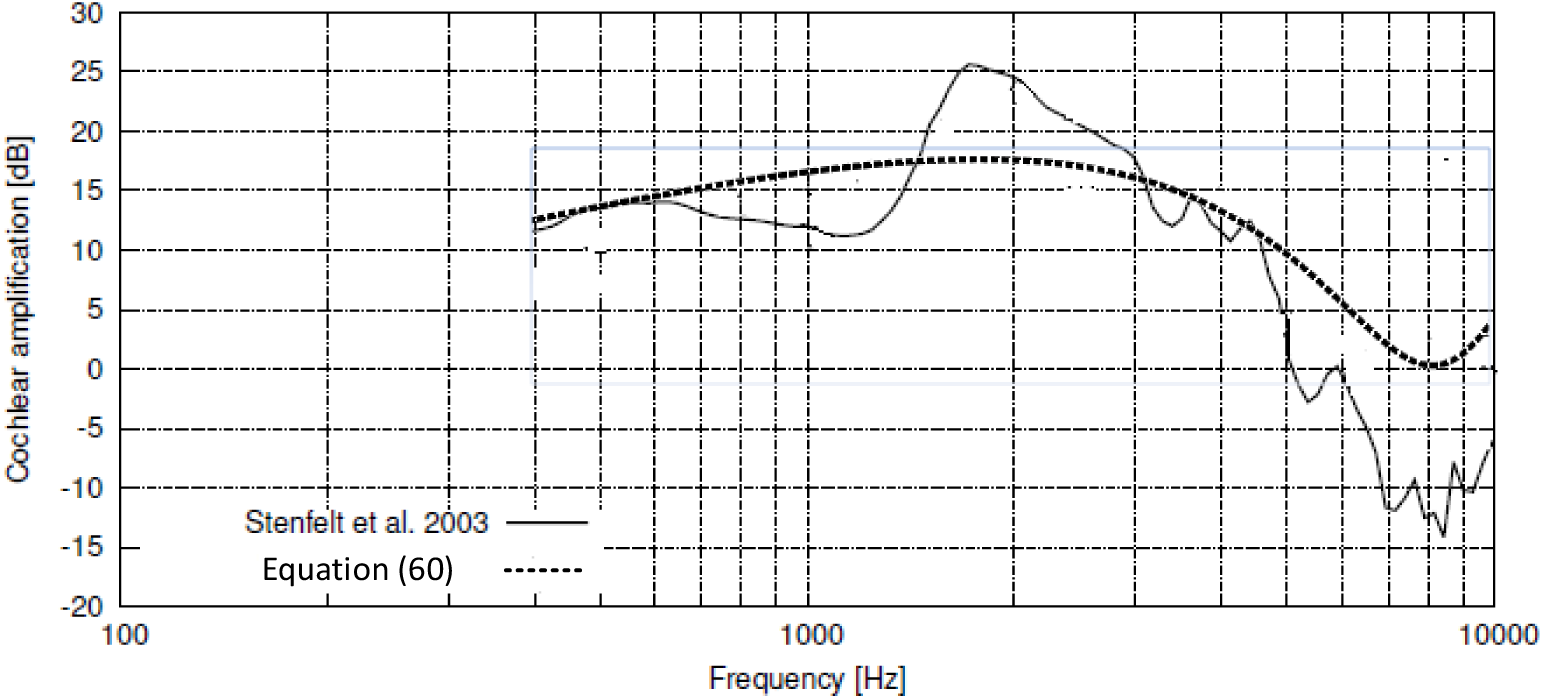
Level of cochlear amplification.

## 4. Conclusions

The shown rules link the values which describe the transfer of the sound within the human ear, when the sound wave with a given intensity and frequency reaches the ear. These values are the amplitude of the stapes footplate, the amplitude of the membrane of the round window, the amplitude of the sound wave in the cochlea and the amplitude of the basilar membrane. The calculations based on these rules give the results close to those received by measurements done by others. The tests were made up of the measurements of amplitudes for the stapes footplate, the round window and the basilar membrane.

Using the given rules, one can do a fast simulation of the ear behavior under an impact of the sound wave. The results should be correct when the sound intensity level lies in the range of 10-120 dB and the frequency in the range of 1-20 kHz. The parameters describing the geometry of the ear can be changed in a wide range. One can use the proposed rules to describe the sound propagation in the ears of the mammals. In that case, one can get more appropriate measurements.

The presented work deals with the process of the ***sound propagation*** in the ear. To describe the process of the ***sound perception***, a proper model of the sound receptors inside the cochlear duct should be given. Here, the cochlear duct is reduced to the basilar membrane. However, the found sound pressure acting on the basilar membrane may be used to predict the motion of the sound receptors.

## Appendix A. Power of sound wave

A plane sound wave runs along the ***x*** axis in a waveguide filled with fluid (liquid or gas) with density ***ρ***. Imagine an element **ℬ** with the cross section area ***F***(***x***) and the width **Δ*x***, cut from the waveguide by the wave fronts at times ***t*** and ***t*** + **Δ*t***. This element has the mass ***ρF***(***x***)**Δ*x*** and moves along the ***x*** axis at the speed of sound ***c*** = **Δ*x***/**Δ*t***.

Consider the energy **Δ*E***(***x, t***) of the particles in such a fluid layer which at the time ***t*** covers with the element **ℬ**. The position and velocity of each of these particles with respect to the wave front at time ***t*** is given by the variables

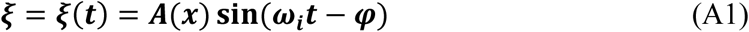

where ***ω***_(***i***)_ is the circular frequency and ***A*** is the amplitude of the sound wave. In addition, ***φ*** is a position of a given particle at the time ***t*** = **0**.

The potential energy **Δ*E***_***p***_(***x, t***) of the above particles is the work of their inertia forces on the way of length ***A***, namely

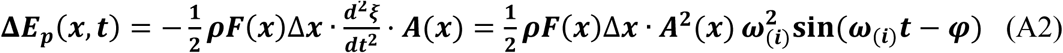

The potential energy **Δ*E***_***k***_(***x, t***) of these particles has the form

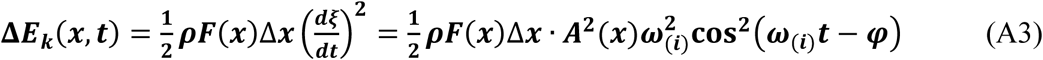

Thus, the total mechanical energy of the fluid layer of the width **Δ*x*** is as follows

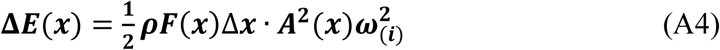

Note that **Δ*E***(***x***) depends on the width **Δ*x*** of the layer, but does not depend on the time ***t***. Moreover, if the wave front in time **Δ*t*** is shifted by **Δ*x***, then at that time the fluid particles obtain energy **Δ*E***(***x***), so in the unit of time they get power

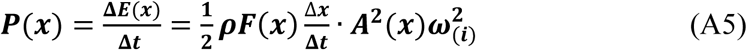

Finally, the power of the sound wave is given by the formula

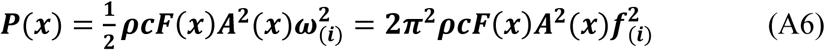

Let ***T***_(***i***)_ = **1/*f***_(***i***)_ = **2*π*/*ω***_(***i***)_ be the period of the running wave, and ***λ***_(***i***)_ it is its length. Consider the stream of fluid cut out from the waveguide by which the wave runs, and whose length is equal to the wavelength ***λ***_(***i***)_. Assume that the particles lying on the initial cross section of the stream have the displacement is ***ξ*** = **0**, at the time ***t*** = **0**. In the fluid stream the wave forms two pairs of self-balanced inertia forces ***N***(***x***). The first pair of the forces causes squeezing of liquid particles in the first half of the wavelength, and the second pair of the forces stretches the other half of the wavelength.

In this way, the ***sound pressure p***(***x***) can be defined as the force ***N***(***x***)acting along the path ***λ***_(***i***)_**/4** and per unit of the waveguide cross section area ***F***(***x***):

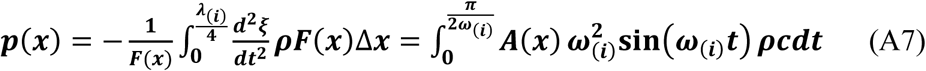

Then, the relation between the sound pressure ***p***_(***i***)_(*x*) and the amplitude of the sound wave ***A***_(***i***)_(*x*) is the following

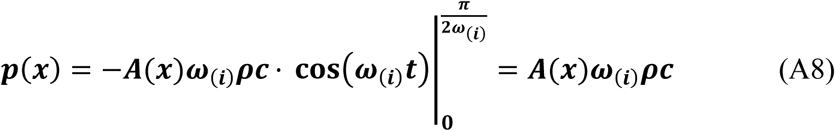

From Eq. (A7), one can express the power of the sound wave ***P***(***x***) by amplitude of the sound pressure ***p***(***x***), namely

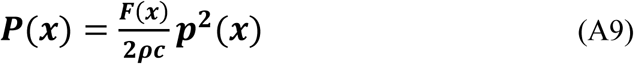

## Appendix B. Effect of basilar membrane waving on sound wave amplitude

Our considerations base on the assumption that the basilar membrane motion has no influence on the sound wave amplitude. Now, we want to get a relation between the amplitudes ***A***_***BM***_(***y***) and ***A***_***C***_(***x***) without this assumption. To do it, let us consider the case when initially the cochlea fluid does not move. Due to vibrations of the stapes footplate, a front of the sound wave begins its run along the ***y***-axis from the oval window to the cochlea apex. The basilar membrane begins to move under the influence of the sound wave pressure

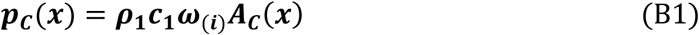

for ***x***_**0**_ ≤ ***x*** = ***y*** + ***x***_**0**_ ≤ ***x***_**0**_ + ***L***_***BM***_. Notice that a conversion of **Δ*A***_***C***_(***y***) to ***A***_***C***_(***x***) in Eqs (45) and 49 gives the relation

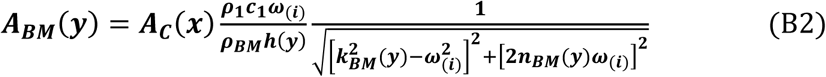

### The principle of the power transmission

Using the principle of power transmission, one can express both ***A***_***BM***_(***y***) and ***A***_***BM***_(***y***) as functions of the stapes footplate amplitude ***A***_***BM***_. However, because the basilar membrane motion may have an influence on the amplitude of the sound wave, instead of Eq. (30), we should to put

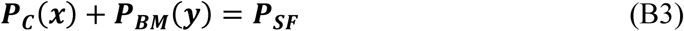

Eqs (27) and (31) show the powers ***P***_***SF***_ and ***P***_***C***_(***x***) as functions of the amplitudes ***P***_***SF***_ and ***A***_***C***_(***x***), respectively. Now, the power of the basilar membrane ***P***_***BM***_(***y***) is derived.

### The power of the basilar membrane *P*_*BM*_(*y*)

Consider a single fiber at the point ***y*** of the basilar membrane of the width **Δ*y***. During **1/4** of the vibration cycle, the fiber bends from the location ***u***_**0**_ = **0** to the position ***u***_**1**_ = ***A***_***BM***_(***y***). The work done by the fiber at time 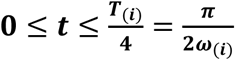 is as follows:

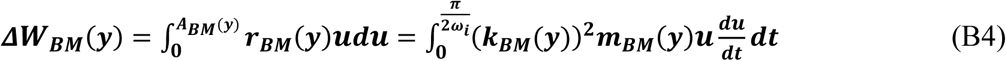

thus

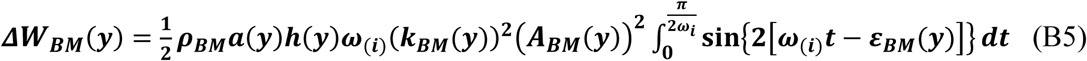

and finally

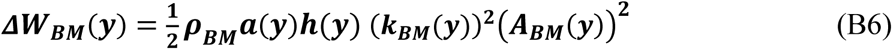

After moving to the position ***u***_**1**_ = ***A***_***BM***_, the fiber at the time ***T***_(***i***)_**/4** ≤ ***t*** ≤ ***T***_(***i***)_**/2** recovers elastic energy and returns to the position ***u***_**0**_ = **0**. Next, the fiber is bent in the opposite direction and come back to the starting position, at the time ***T***_(***i***)_**/2** ≤ ***t*** ≤ ***T***_(***i***)_, doing the work given by Eq. (B6). Concluding, the power of the fiber ***ΔP***_***BM***_(***y***) is the work **2*ΔW***_***BM***_(***y***) done at time ***T***_***i***_, namely

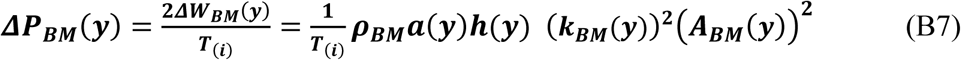

Now, one can describe the power of this part of the basement membrane over which the sound wave front has passed.

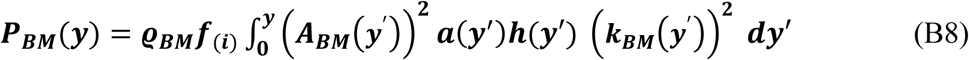

### The interaction between the basilar membrane and the sound wave

From Eq. (B3) yields the following integral equation.

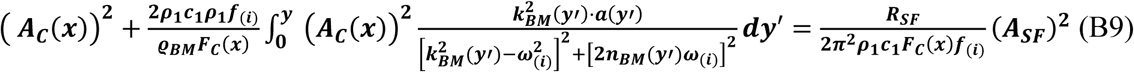

It can be changed into a differential equation for ***A***_***C***_(***x***) in which ***y*** = ***x*** − ***x***_**0**_

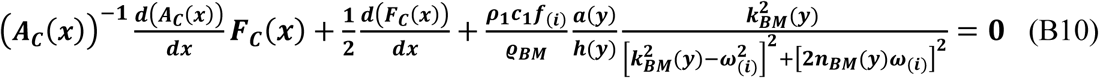

with a boundary condition:

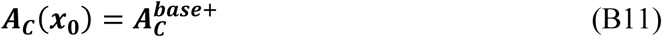

Eq. (B10) may be shown in the form:

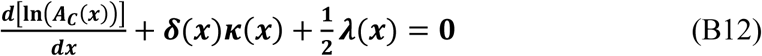

where 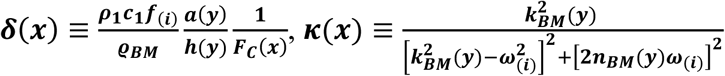 and 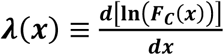.

A general solution of Eq. (B12) is as follows.

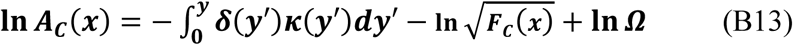

One can get the constant ***Ω*** from the boundary condition (B11):

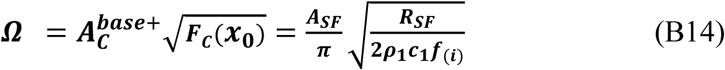

Finally, we have

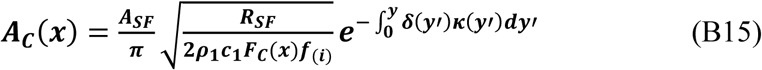

Comparing Eqs (32) and (B15), one can see that the influence of the basilar membrane motion on the amplitude ***A***_***C***_(***x***) of the sound wave is due to the multiplier 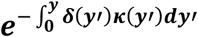. One can show that at the cochlea apex, where ***x*** = ***L***_***C***_, for the considered range of frequencies, there is

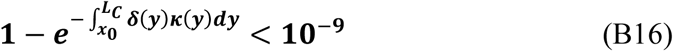

Thus, one can state that the basilar membrane motion has no influence on the amplitude of the sound wave. Concluding, Eq. (B15) may be written in the form

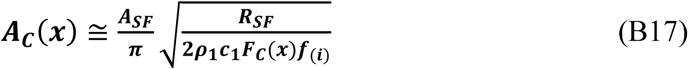

## Appendix C. List of symbols and calculation data

Below is a list of the used parameters, their units, fixed values and basic relations, in the order in which they appear. If a numerical value of some of these parameters is not given, then this value can be got on the base of the other parameters.

### 1° Outer ear

***β*** [*dB*] **−** level of sound intensity,

***f***_(***i***)_ [*1/s*] – sound frequency.

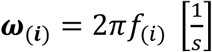 – sound angular frequency,

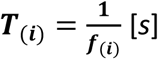 – sound wave period,

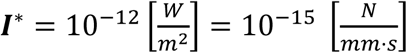 **−** reference sound intensity,

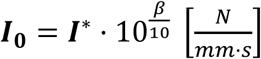 – sound intensity received by outer ear,

***F***_**0**_ = 35 [*mm*^2^] – cross-sectional area of entrance to auditory canal,

***F***_***TM***_ = 9.85 [*mm*^2^] – cross-sectional area of tympanic membrane base,

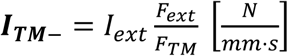 – sound intensity just before tympanic membrane

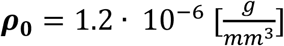 – air density,

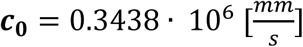 – sound velocity in air (at 20°C),

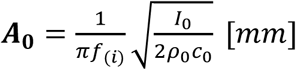 **−** sound amplitude at ear canal inlet,

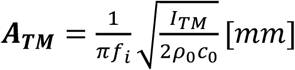 – sound amplitude just before tympanic membrane,

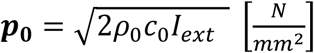 – sound pressure amplitude at ear canal inlet,

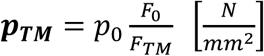 – sound pressure amplitude just before tympanic membrane.

### 2° Middle ear

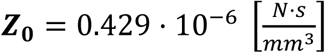 – wave resistance of air,

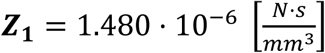 – wave resistance of cochlear fluid,

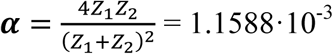 – absorption coefficient of sound wave,

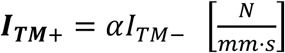 – sound intensity transmitted to middle ear,

***N***_***TM***–_ = *p*_*TM*_*F*_*TM*_ [*N*] – maximal force acting on tympanic membrane,

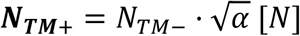 – maximal force acting on malleus,

***γ*** = 1.3 – malleus-incus lever transmission,

***N***_***SF***_ = ***γ****N*_***TM***+_ [*N*] – maximal force acting on stapes footplate,

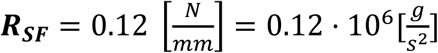 – stiffness of annular ligament of stapes footplate,

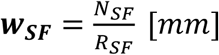 – static amplitude of stapes footplate,

***M***_***SF***_ = 3.04 · 10^−3^ [*g*] – mass of stapes,

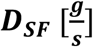 – damping of stapes footplate,

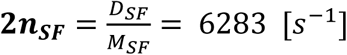 – damping coefficient of stapes footplate,

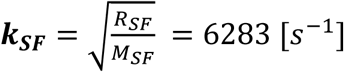 – resonant angular frequency of stapes footplate,

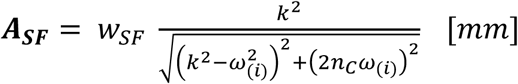 – dynamic amplitude of stapes footplate,

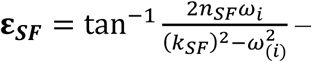 **−** phase shift of stapes footplate.

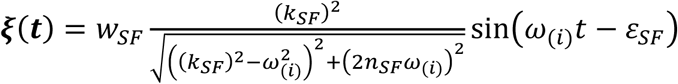 **−** stapes footplate displacement.

### 3° Round window

***F***_***SF***_ = 2.5[*mm*^2^] – active area of stapes footplate,

***r***_***RW***_ = 0.6 [*mm*] – radius of round window membrane,

***F***_***RW***_ = ***πr***_***RW***_^2^ = 1.13 [*mm*^2^] –area of round window membrane,

***w***_***RW***_ [*mm*] **−** static amplitude of round window membrane,

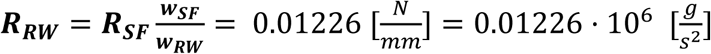 – stiffness of round window membrane,

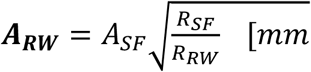 **−** amplitude of round window membrane.

### 4° Cochlea

***L***_***C***_ = 34.5 [*mm*] – length of cochlea axis,

***x*** [*mm*] (0 ≤ ***x*** ≤ 2***L***_***C***_) – distance along cochlea axis from oval window,

***b***(***x***) [*mm*] – width of cochlea,

***b***(***x***_**0**_) = 4.3 [*mm*] – width of cochlea at the base of basilar membrane,

***b***(***L***_***C***_) = 1.7 [*mm*] – width of cochlea at the apex,

***d*′**(***x***) [*mm*] – height of scala vestibuli,

***d*′**(***x***_**0**_) = 2.0 [*mm*] – height of scala vestibuli at the base of basilar membrane,

***d*′**(***L***_***C***_) = 1.0 [*mm*] – height of scala vestibuli at the apex,

***d*″**(***x***) [*mm*] – height of scala tympani,

***d*″**(***x***_**0**_) = 1.5 [*mm*] – height of scala tympani at the base of basilar membrane,

***d*″**(***L***_***C***_) = 0.7 [*mm*] – height of scala tympani at the apex,

***F***_***C***_(***x***) [*mm*^2^] – area of cochlea cross section,

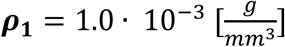 – cochlea fluid density,

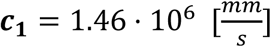 – sound velocity in cochlea fluid,

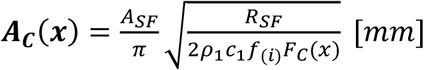 – sound wave amplitude in cochlea,

***p***(***x***)– sound wave pressure in cochlea,

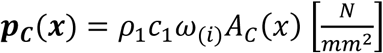 – sound pressure amplitude in cochlea,

### 4° Basilar membrane

***L***_***BM***_ = 31.9 [*mm*] – length of basilar membrane,

***x***_**0**_ = 2.6 [*mm*] **−** distance basilar membrane base from oval window,

***a***(***y***) [*mm*] (***x***_0_ ≤ ***x*** ≤ 2***L***_***BM***_) – width of basilar membrane,

***a***(**0**) = 0.1 [*mm*] – width of basilar membrane at the base,

***a***(***L***_***BM***_) = 0.5 [*mm*] – width of basilar membrane at the apex,

***h***(***y***) [*mm*] (0 ≤ ***x*** ≤ 2***L***_***C***_) – height of basilar membrane,

***h***(**0**) = 0.075 [*mm*] – height of basilar membrane at the base,

***h***(***L***_***BM***_) = 0.025 [*mm*] – height of basilar membrane at the apex,

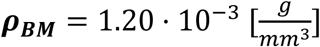 – basilar membrane density,

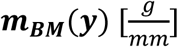 – normalized mass of basilar membrane fiber (per unit of fiber width),

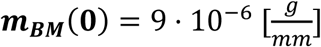 **−** normalized mass of basilar membrane at the base,

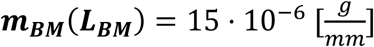 **−** normalized mass of basilar membrane at the apex,

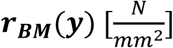 – normalized stiffness of basilar membrane fiber (per unit of fiber width),

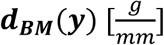 – normalized damping of basilar membrane fiber (per unit of fiber width),

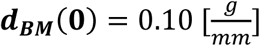 – normalized damping of basilar membrane at the base,

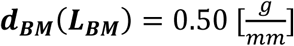 – normalized damping of basilar membrane at the apex,

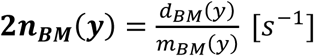 – damping coefficient of basilar membrane fiber,

**2*n***_***BM***_(***y***) = 11111 [*s*^−1^] – damping coefficient of basilar membrane at the base,

**2*n***_***BM***_(***y***) = 3.3333 [*s*^−1^] – damping coefficient of basilar membrane at the apex,

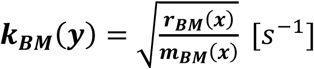 – resonant angular frequency of basilar membrane fiber,

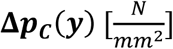 – differential pressure acting on basilar membrane,

**Δ*A***_***C***_(***y***) [*mm*] **−** differential sound wave amplitude,

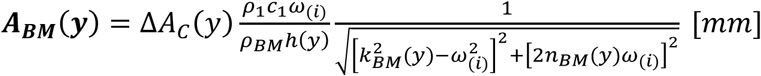 – BM amplitude,

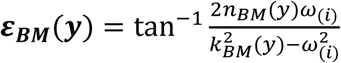 **−** phase shift of basilar membrane,

***u***(***y, t***) = *A*_*BM*_(*y*)sin[*ω*_(*i*)_*t* − *ε*_*BM*_(*y*)] [*mm*] **−** basilar membrane displacement.

